## Supplementary File for "Mesolimbic dopamine projections mediate cue-motivated reward seeking but not reward retrieval"

**Table 1. Generalized linear mixed-effects model output for the analysis of total lever presses during PIT trials in rats expressing hM4Di or mCherry (Experiment 2).**

|  | *t*(240) | *P* | *b* | 95% CI |
| --- | --- | --- | --- | --- |
| Intercept | 23.01 | < .001 | 2.66 | [2.43, 2.89] |
| Group | -0.10 | .920 | -0.01 | [-0.24, 0.22] |
| Drug | -1.08 | .279 | -0.05 | [-0.15, 0.04] |
| CS Period | 5.07 | < .001 | 0.21 | [0.13, 0.29] |
| CS Type | 2.57 | .011 | 0.11 | [0.03, 0.19] |
| Group * Drug | -0.43 | .670 | -0.02 | [-0.12, 0.07] |
| Group * CS Period | -0.65 | .517 | -0.03 | [-0.11, 0.05] |
| Drug * CS Period | -2.62 | .009 | -0.04 | [-0.07, -0.01] |
| Group * CS Type | 0.44 | .659 | 0.02 | [-0.07, 0.10] |
| Drug * CS Type | -0.49 | .627 | -0.01 | [-0.04, 0.02] |
| CS Period * CS Type | 6.24 | < .001 | 0.10 | [0.07, 0.13] |
| Group * Drug * CS Period | -0.42 | .677 | -0.01 | [-0.04, 0.02] |
| Group * Drug * CS Type | 2.76 | .006 | 0.04 | [0.01, 0.07] |
| Group * CS Period * CS Type | 0.26 | .796 | 0.004 | [-0.03, 0.03] |
| Drug * CS Period * CS Type | -2.71 | .007 | -0.04 | [-0.07, -0.01] |
| Group * Drug * CS Period * CS Type | -3.14 | .002 | -0.05 | [-0.08, -0.02] |

Note: Categorical variables were effect coded with mCherry/hM4Di (Group) as ‑1/+1, Vehicle/CNO (Drug) as ‑1/+1, Pre/CS (CS Period) as ‑1/+1, and Unpaired/Paired (CS Type) as ‑1/+1. Analysis included 256 data points (df = 240). *b* = unstandardized regression coefficient; CI = confidence interval; CS = conditioned stimulus. See Figure 3B for data presentation.

**Table 2. Generalized linear mixed-effects model output for the analysis of the proportion of presses that were followed by a food-cup approach during PIT trials in rats expressing hM4Di or mCherry (Experiment 2).**

|  | *t*(176) | *P* | *b* | 95% CI |
| --- | --- | --- | --- | --- |
| Intercept | 26.12 | < .001 | 0.46 | [0.43, 0.50] |
| Group | -0.42 | .676 | -0.01 | [-0.04, 0.03] |
| Drug | -2.37 | .019 | -0.03 | [-0.05, -0.01] |
| CS Period (CS-) | -4.33 | < .001 | -0.08 | [-0.11, -0.04] |
| CS Period (CS+) | 7.64 | < .001 | 0.14 | [0.10, 0.17] |
| Group * Drug | 0.02 | .984 | 0.0003 | [-0.02, 0.03] |
| Group * CS Period (CS-) | 0.06 | .953 | 0.001 | [-0.03, 0.04] |
| Group * CS Period (CS+) | -0.01 | .993 | -0.0002 | [-0.04, 0.03] |
| Drug * CS Period (CS-) | -0.84 | .403 | -0.02 | [-0.05, 0.02] |
| Drug * CS Period (CS+) | 2.10 | .037 | 0.04 | [0.002, 0.07] |
| Group * Drug * CS Period (CS-) | 0.32 | .748 | 0.01 | [-0.03, 0.04] |
| Group * Drug * CS Period (CS+) | -0.21 | .835 | -0.004 | [-0.04, 0.03] |

Note: For analysis, data were square-root transformed because their distribution was positive skewed. Categorical variables were effect coded with mCherry/hM4Di (Group) as ‑1/+1, and Vehicle/CNO (Drug) as ‑1/+1. The categorical variable CS Period had 3 levels (Pre, CS-, CS+), and the Pre-CS period served as the reference level. Analysis included 188 data points (df = 176). For the main effect of and interactions involving CS Period, *t*-values refer to the simple effects of each level of CS Period versus the population mean. The overall significance of the main effect and interaction are reported in the main text as *F*-values, as described in the Supplementary Methods. *b* = unstandardized regression coefficient; CI = confidence interval; CS = conditioned stimulus. See Figure 3D for data presentation.

**Table 3. Generalized linear mixed-effects model output for the analysis of total lever presses during PIT trials in rats expressing the inhibitory DREADD hM4Di and receiving CNO or vehicle microinfusions in either the NAc or mPFC (Experiment 3a).**

|  | *t*(240) | *P* | *b* | 95% CI |
| --- | --- | --- | --- | --- |
| Intercept | 14.30 | < .001 | 1.81 | [1.56, 2.05] |
| Drug | -1.13 | .260 | -0.09 | [-0.25, 0.07] |
| CS Period | 8.84 | < .001 | 0.47 | [0.37, 0.58] |
| CS Type | 4.26 | < .001 | 0.20 | [0.11, 0.29] |
| Site | 0.13 | .897 | 0.02 | [-0.23, 0.27] |
| Drug * CS Period | -0.13 | .894 | -0.003 | [-0.05, 0.05] |
| Drug * CS Type | -4.04 | < .001 | -0.10 | [-0.16, -0.05] |
| CS Period * CS Type | 9.41 | < .001 | 0.24 | [0.19, 0.29] |
| Drug * Site | 0.62 | .536 | 0.05 | [-0.11, 0.21] |
| CS Period * Site | 0.68 | .500 | 0.04 | [-0.07, 0.14] |
| CS Type * Site | -2.09 | .038 | -0.10 | [-0.19, -0.01] |
| Drug * CS Period * CS Type | 0.72 | .472 | 0.02 | [-0.03, 0.07] |
| Drug * CS Period * Site | -3.06 | .002 | -0.08 | [-0.13, -0.03] |
| Drug * CS Type * Site | 1.07 | .285 | 0.03 | [-0.02, 0.08] |
| CS Period * CS Type * Site | -0.76 | .451 | -0.02 | [-0.07, 0.03] |
| Drug * CS Period * CS Type * Site | -2.99 | .003 | -0.08 | [-0.13, -0.03] |

Note: Categorical variables were effect coded with Vehicle/CNO (Drug) as ‑1/+1, Pre/CS (CS Period) as ‑1/+1, Unpaired/Paired (CS Type) as ‑1/+1, and mPFC/NAc (Site) as -1/+1. Analysis included 256 data points (df = 240). *b* = unstandardized regression coefficient; CI = confidence interval; CS = conditioned stimulus. See Figure 4C for data presentation.

**Table 4. Generalized linear mixed-effects model output for the analysis of the proportion of presses followed by a food-cup approach during PIT trials in rats expressing the inhibitory DREADD hM4Di and receiving CNO or vehicle microinfusions in either the NAc or mPFC (Experiment 3a).**

|  | *t*(139) | *P* | *b* | 95% CI |
| --- | --- | --- | --- | --- |
| Intercept | 12.35 | < .001 | 0.42 | [0.35, 0.49] |
| Drug | -1.92 | .057 | -0.03 | [-0.07, 0.001] |
| Site | -0.40 | .687 | -0.01 | [-0.08, 0.05] |
| CS Period (CS-) | -5.55 | < .001 | -0.14 | [-0.18, -0.09] |
| CS Period (CS+) | 7.23 | < .001 | 0.17 | [0.13, 0.22] |
| Drug * Site | 0.04 | .969 | 0.001 | [-0.04, 0.04] |
| Drug * CS Period (CS-) | 0.11 | .911 | 0.003 | [-0.05, 0.05] |
| Drug * CS Period (CS+) | 1.16 | .250 | 0.03 | [-0.02, 0.08] |
| Site * CS Period (CS-) | -0.87 | .385 | -0.02 | [-0.07, 0.03] |
| Site * CS Period (CS+) | 1.07 | .288 | 0.03 | [-0.02, 0.07] |
| Drug * Site * CS Period (CS-) | -1.23 | .222 | -0.03 | [-0.08, 0.02] |
| Drug * Site * CS Period (CS+) | 1.69 | .093 | 0.04 | [-0.01, 0.09] |

Note: For analysis, data were square-root transformed because their distribution was positive skewed. Categorical variables were effect coded with Vehicle/CNO (Drug) as ‑1/+1, and mPFC/NAc (Site) as -1/+1. The categorical variable CS Period had 3 levels (Pre, CS-, CS+), and the Pre-CS period served as the reference level. Analysis included 151 data points (df = 139). *b* = unstandardized regression coefficient; CI = confidence interval; CS = conditioned stimulus. See Figure 4E for data presentation.

**Table 5. Generalized linear mixed-effects model output for the analysis of total lever presses during reward devaluation (Nonreinforced phase; Experiment 4).**

|  | *t*(148) | *P* | *b* | 95% CI |
| --- | --- | --- | --- | --- |
| Intercept | 22.79 | < .001 | 2.59 | [2.36, 2.81] |
| Group | 0.48 | .634 | 0.05 | [-0.17, 0.28] |
| Drug | -0.19 | .849 | -0.01 | [-0.09, 0.08] |
| Lever | -5.41 | < .001 | -0.67 | [-0.91, -0.42] |
| Group * Drug | 0.15 | .879 | 0.01 | [-0.08, 0.09] |
| Group * Lever | 0.84 | .404 | 0.10 | [-0.14, 0.35] |
| Drug * Lever | 1.46 | .146 | 0.03 | [-0.01, 0.08] |
| Group * Drug * Lever | 0.54 | .591 | 0.01 | [-0.03, 0.06] |

Note: Categorical variables were effect coded with mCherry/hM4Di (Group) as ‑1/+1, Vehicle/CNO (Drug) as ‑1/+1, and Valued/Devalued (Lever) as ‑1/+1. Analysis included 156 data points (df = 148). *b* = unstandardized regression coefficient; CI = confidence interval. See Figure 5B for data presentation.

**Table 6. Generalized linear mixed-effects model output for the analysis of the proportion of presses followed by a food-cup approach during reward devaluation (Nonreinforced phase; Experiment 4).**

|  | *t*(139) | *P* | *b* | 95% CI |
| --- | --- | --- | --- | --- |
| Intercept | 17.51 | < .001 | 0.54 | [0.48, 0.60] |
| Group | -0.45 | .652 | -0.01 | [-0.08, 0.05] |
| Drug | 0.54 | .591 | 0.01 | [-0.03, 0.04] |
| Lever | 2.08 | .040 | 0.04 | [0.002, 0.07] |
| Group * Drug | -0.15 | .881 | -0.003 | [-0.04, 0.03] |
| Group * Lever | 0.41 | .680 | 0.01 | [-0.03, 0.04] |
| Drug * Lever | 1.61 | .109 | 0.03 | [-0.01, 0.06] |
| Group * Drug * Lever | -0.56 | .578 | -0.01 | [-0.04, 0.02] |

Note: For analysis, data were square-root transformed because their distribution was positive skewed. Categorical variables were effect coded with mCherry/hM4Di (Group) as ‑1/+1, Vehicle/CNO (Drug) as ‑1/+1, and Valued/Devalued (Lever) as ‑1/+1. Analysis included 147 data points (df = 139). *b* = unstandardized regression coefficient; CI = confidence interval. See Figure 5C for data presentation.

**Table 7. Generalized linear mixed-effects model output for the analysis of lever presses that were performed with or without a subsequent food-cup approach response during reward devaluation (Nonreinforced phase; Experiment 4).**

|  | *t*(296) | *P* | *b* | 95% CI |
| --- | --- | --- | --- | --- |
| Intercept | 15.204 | < .001 | 1.71 | [1.49, 1.92] |
| Group | 0.56 | .577 | 0.06 | [-0.16, 0.28] |
| Drug | -0.14 | .885 | -0.01 | [-0.09, 0.08] |
| Press Type | 8.10 | < .001 | 0.55 | [0.42, 0.68] |
| Lever | -4.93 | < .001 | -0.57 | [-0.80, -0.34] |
| Group * Drug | 0.25 | .799 | 0.01 | [-0.08, 0.10] |
| Group * Press Type | 0.56 | .576 | 0.04 | [-0.10, 0.17] |
| Drug * Press Type | 0.18 | .859 | <0.01 | [-0.04, 0.05] |
| Group * Lever | 0.75 | .455 | 0.09 | [-0.14, 0.31] |
| Drug * Lever | 0.90 | .369 | 0.02 | [-0.03, 0.07] |
| Press Type * Lever | -7.92 | <.001 | -0.21 | [-0.26, -0.16] |
| Group * Drug * Press Type | -0.22 | .822 | -0.01 | [-0.05, 0.04] |
| Group * Drug * Lever | 0.03 | .975 | <0.01 | [-0.05, 0.05] |
| Group * Press Type * Lever | 2.07 | .040 | 0.05 | [>0.00, 0.11] |
| Drug * Press Type * Lever | 0.99 | .320 | 0.02 | [-0.02, 0.07] |
| Group * Drug * Press Type * Lever | 0.84 | .401 | 0.02 | [-0.03, 0.07] |

Note: Categorical variables were effect coded with mCherry/hM4Di (Group) as ‑1/+1, Vehicle/CNO (Drug) as ‑1/+1, Presses With/Without Approach (Press Type) as -1/+1, and Valued/Devalued (Lever) as ‑1/+1. Analysis included 312 data points (df = 296). *b* = unstandardized regression coefficient; CI = confidence interval. See Figure 5D for data presentation.
